# Regional context for balancing sagebrush- and woodland-dependent songbird needs with targeted pinyon-juniper management in the sagebrush biome

**DOI:** 10.1101/2022.05.03.490495

**Authors:** Jason D. Tack, Joseph T. Smith, Kevin E. Doherty, Patrick J. Donnelly, Jeremy D. Maestas, Brady W. Allred, Jason Reinhardt, Scott L. Morford, David E. Naugle

## Abstract

Tree expansion among historic grassland and shrubland systems is a global phenomenon, which results in dramatic influences on ecosystem processes and wildlife populations. In the western US, pinyon-juniper woodlands have expanded by as much as six-fold among sagebrush steppe landscapes since the late nineteenth century, with demonstrated negative impacts to the behavior, demography, and population dynamics of species that rely on intact sagebrush rangelands. Notably, greater sage-grouse (*Centrocercus urophasianus*) are unable to tolerate even low conifer cover, which can result in population declines and local extirpation. Removing expanding conifer cover has been demonstrated to increase sage grouse population growth rates and sagebrush-obligate songbird abundance. However, advances in restoring sagebrush habitats have been met with concern about unintended impacts to species that rely on conifer woodlands, notably the pinyon jay (*Gymnorhinus cyanocephalus*) whose population declines are distinctive among birds breeding in pinyon-juniper woodlands. We modeled indices to abundance in relation to multi-scale habitat features for nine songbirds reliant on both sagebrush and pinyon-juniper woodlands for breeding. Findings demonstrate that targeted sage grouse habitat restoration under the Sage Grouse Initiative is not at odds with protection of pinyon jay populations. Rather, conifer management has largely occurred in the northern sagebrush ecosystem where models suggest that past cuts likely benefit Brewer’s sparrow and sage thrasher while avoiding pinyon jay habitat. Extending our spatial modeling further south beyond the sagebrush biome could better equip conservationists with more comprehensive decision-support, particularly where pinyon jays face additional pressures of drought-induced tree mortality.

## Introduction

Native tree species are expanding into shrublands and grasslands globally at an alarming rate, increasing from 40-600% in distribution across every continent except Antarctica (Nackley et al., 2017). Resulting shifts in vegetation structure and composition are affecting a broad suite of ecosystem services and values, including wildlife species of conservation concern (Baruch-Mordo et al., 2013; Fuhlendorf et al., 2017, 2002). In North America, pinyon-juniper woodlands, composed of both juniper (*Juniperus spp.*) and pinyon pine (*Pinus spp.*; hereafter collectively referred to as conifer), are among the most dominant vegetation types across the intermountain western United States, supporting critical biodiversity, ecosystem services, and economic potential (Romme et al., 2009). Since European settlement, the distribution of these conifer species has expanded between two- and six-fold, likely due to the compounding effects of historic high-intensity grazing, subsequent increases in natural fire return intervals that limited woodland establishment, and favorable climatic conditions that helped tree growth proliferate among sagebrush (*Artemisia spp.*) systems (Miller et al., 2019). Some 90% of pinyon-juniper expansion has occurred in sagebrush ecosystems (Miller et al., 2011), leading to a loss of sagebrush and herbaceous vegetation (Roundy et al., 2014) and associated specialist wildlife species (Baruch-Mordo et al., 2013; Rickart et al., 2008). As a result, management to remove conifers from former shrublands has been adopted as a widespread conservation practice to mitigate negative ecosystem impacts over the past decade (Miller et al., 2017).

Central to the proliferation of recent restoration efforts is the conservation of sagebrush-obligate wildlife under the umbrella of greater (*Centrocercus urophasianus*) and Gunnison sage-grouse (*C. minimus*; hereafter collectively referred to as sage grouse; Doherty et al., 2018; Miller et al., 2017). Sage grouse are particularly vulnerable to conifer expansion (Baruch-Mordo et al., 2013). Conifer presence reduces the quality of sage grouse habitat through both behavioral avoidance by nesting females (Severson et al., 2017a) and the demographic consequences of reduced nest (Severson et al., 2017b), brood (Sandford et al., 2017), and female survival (Coates et al., 2017). Experimental research among conifer removal projects has demonstrated that sage grouse quickly return to restored habitats, with subsequent increases in nest, brood, and female survival in treated areas (Sandford et al., 2017; Severson et al., 2017b, 2017c). Ultimately, restoration of habitats through conifer management is translating into measurable population benefits at watershed scales, accounting for a 12% increase in population growth rates compared to control areas in southern Oregon (Olsen et al., 2021). Efficacy of accelerating investments in large-scale restoration efforts via conifer removal was one key factor in obviating the need for an Endangered listing status for sage grouse (US Fish and Wildlife Service 2015), which continues to be a primary management practice for voluntary conservation of sagebrush habitats (Natural Resources Conservation Service 2021).

Benefits from conifer removal targeted for sage grouse likely accrue for other sagebrush-obligate species, though few studies have actually measured resulting benefits of management across taxa (Bombaci and Pejchar, 2016; Zeller et al., 2021). Conifer removal projects for sage grouse have had a high congruence with the predicted distributions of certain sagebrush-obligate songbirds (Donnelly et al., 2017), and past management has resulted in local increases in abundances of shrubland species including Brewer’s sparrow (*Spizella breweri*) and green-tailed towhee (*Pipilo chlorurus*; Holmes et al., 2017). Conversely, the potential for unintended negative impacts to species reliant on conifer woodlands remains a pervasive question, especially for non-target songbirds species of conservation concern (Boone et al., 2018; Zeller et al., 2021).

Among songbirds in the western US, those reliant on pinyon-juniper or sagebrush for breeding habitat have largely demonstrated contrasting population trends over the past 50 years that is consistent with an expanding footprint of conifer among sagebrush habitats (Table 1). Brewer’s sparrow, green-tailed towhee, and sage thrasher (*Oreoscoptes montanus*), all reliant on unfragmented sagebrush habitats for breeding, have experienced population declines; whereas woodland songbirds including ash-throated flycatcher (*Myiarchus cinerascens*), gray flycatcher (*Empidonax wrightii*), gray vireo (*Vireo vicinior*), and juniper titmouse (*Baeolophus ridgwayi*) exhibit stable to increasing populations (Sauer et al., 2017; Table 1). The one notable exception among woodland-reliant species is the pinyon jay (*Gymnorhinus cyanocephalus*), which depends on a mutualistic relationship with conifer nut production for their survival and reproduction (Ligon, 1978). Pinyon jays have declined more severely since 1968 than any other land bird inhabiting sagebrush-associated landscapes (Boone et al., 2018; Sauer et al., 2017), with concern culminating in the US Fish and Wildlife Service being petitioned to list the pinyon jay as Threatened or Endangered under the Endangered Species Act (https://defenders.org/sites/default/files/inline-files/2022.4.25_FWS_Listing%20petition_Pinyon%20Jay.pdf; accessed 3 May, 2022).

**Table 1.** Songbird species used to develop species distribution models and resulting spatial predictions from Breeding Bird Survey (BBS) Data. Sagebrush- and woodland-dependent species were considered for modeling, and we used BBS survey-wide estimated trends from 1966-2015 to identify if populations were increasing, decreasing based on direction of 80% of the credible intervals ([CI] *denotes CI overlapping 0; Sauer et al., 2017). We only modeled species if they were detected at least 5 times within Commission for Environmental Cooperation Level 3 ecoregions from 2011-2016 (supplemental Fig. 1 for map and corresponding ecoregion names).

| Species | Habitat | BBS Trend | Ecoregions |
| --- | --- | --- | --- |
| Brewer's Sparrow | Sagebrush | Declining; -1.01 (-1.89, -0.22) | 1, 2, 4, 5, 6, 7, 8, 9, 10, 11, 13, 14, 15, 16, 17, 18, 19, 20, 21, 22, 23, 24 |
| Green-tailed Towhee | Sagebrush | Declining* ; -0.31 (-0.83, 0.19) | 1, 2, 4, 5, 6, 9, 11, 13, 16, 18, 19, 20, 21, 23, 24 |
| Sagebrush Sparrow | Sagebrush | Increasing* ; 0.43 (-3.51, 4.51) | 1, 2, 5, 6, 8, 13, 16, 18, 20, 21, 23, 24 |
| Sage Thrasher | Sagebrush | Declining; -1.20 (-1.93, -0.47) | 1, 2, 5, 6, 8, 9, 11, 13, 16, 17, 18, 19, 20, 21, 23, 24 |
| Ash-throated Flycatcher | Woodland | Increasing; 1.10 (0.68, 1.52) | 1, 2, 3, 4, 5, 6, 8, 9, 12, 14, 16, 19, 21, 22, 23, 24 |
| Gray Flycatcher | Woodland | Increasing; 2.43 (1.45, 3.51) | 1, 2, 5, 6, 7, 8, 9, 13, 14, 16, 19, 21, 23, 24 |
| Gray Vireo | Woodland | Increasing; 2.10 (-0.28, 4.24) | 1, 5, 6, 14, 21, 23, 24 |
| Juniper Titmouse | Woodland | Increasing* ; 0.28 (-1.14, 1.54) | 1, 5, 6, 9, 14, 16, 21, 22, 23, 24 |
| Pinyon Jay | Woodland | Declining; -3.69 (-5.08, -2.37) | 1, 2, 5, 6, 9, 13, 14, 16, 18, 19, 21, 23, 24 |

Mechanisms underlying pinyon jay population declines within sagebrush ecosystems are unknown. Factors hypothesized to contribute to declines include climate-mediated declines in pinyon pine seed production, intentional pinyon-juniper removal, tree die-off, wildfire, and drought; and transition of the preferred heterogeneous pinyon pine and sagebrush stands to persistent woodlands by a process known as “infill” (Boone et al., 2018). Following the infill of mixed sagebrush and conifer stands, individual trees have reduced seed productivity, thus conifer infill may be analogously detrimental to pinyon jay as encroachment is to sagebrush-obligate wildlife like the sage grouse (Fig. 1). Ultimately, improved spatial planning products for both sagebrush-and woodland-obligate birds of conservation concern are needed to enable informed decisions about potential impacts of ongoing management, and foster a holistic approach to multiple species management along the shrubland-to-woodland continuum (Maestas et al. 2021).

**Figure 1.**
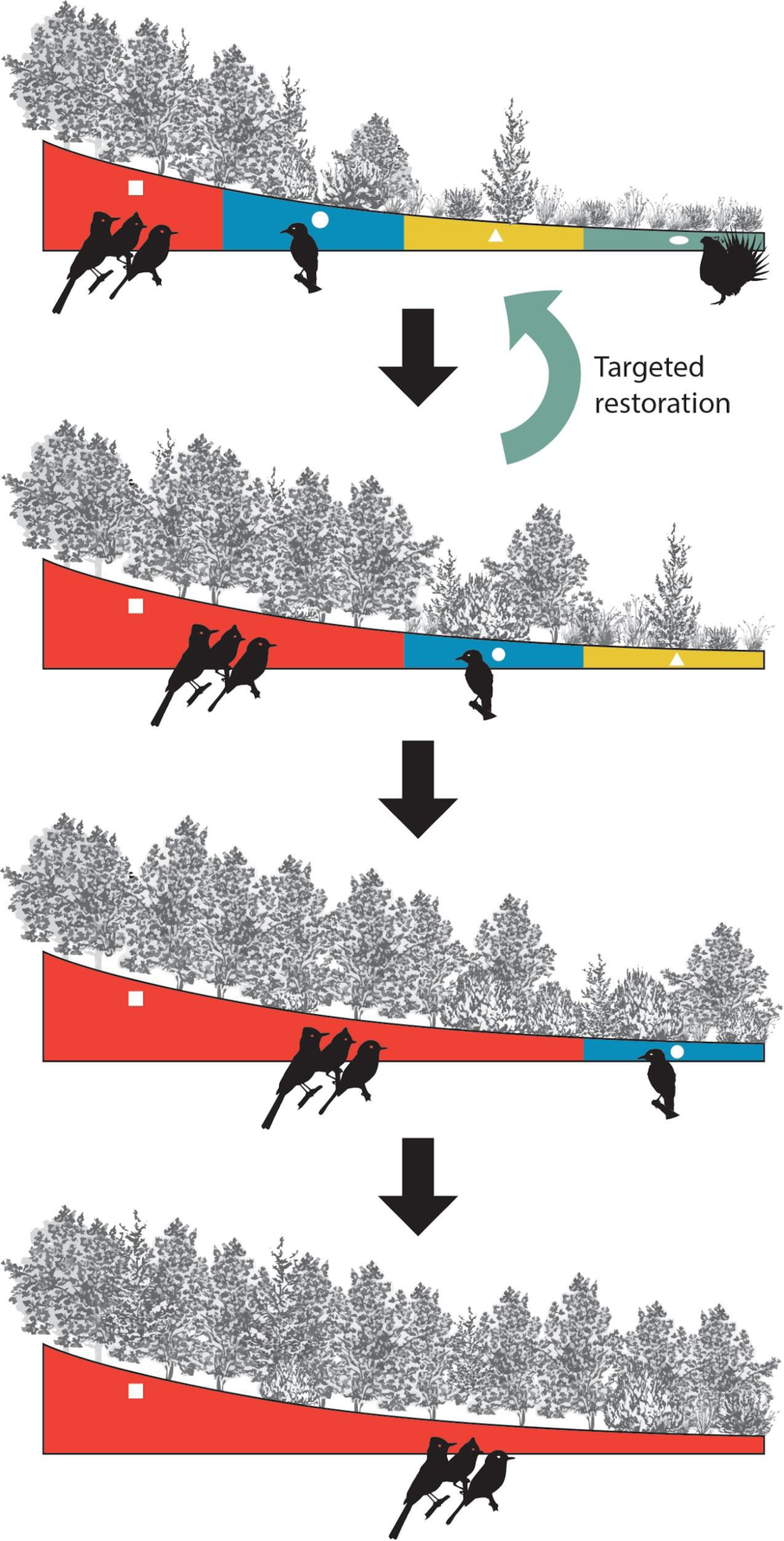
Hypothetical illustration depicting how conifer expansion and infill may impact habitats for both sagebrush-obligate and woodland-reliant birds. The top panel shows a landscape that supports a diversity of bird species partitioned by different ecological sites: persistent woodlands supporting dense forest birds (red/square), heterogeneous woodlands supporting birds reliant on more open stands, such as, pinyon jay (blue/circle), and sagebrush shrublands supporting obligate birds, such as, sage grouse where encroaching conifers are targeted for restoration (yellow/triangle to green/oval). Remaining panels depict shifting habitat niches as conifer expansion and infill, without intervention, displace species like pinyon jay and sage grouse that rely on mixed woodlands or treeless shrublands. BBS trends lend support to this hypothetical scenario as both pinyon jay and sagebrush-obligates have been been in decline, while other songbirds reliant on persistent pinyon-juniper woodlands have been increasing (Table 1).

We used Breeding Bird Survey (BBS) data to develop spatial models predictive of the relative abundance of both sagebrush and woodland obligate songbirds inhabiting sagebrush steppe. Maps were developed to: 1) depict relative species distributions spatially, 2) evaluate recent conifer removal for sage grouse in relation to predicted songbird distributions, and 3) help guide spatial targeting of future conservation actions. Applying spatial models to BBS data provides an effective tool to learn about large-scale distribution of breeding birds (Niemuth et al., 2017). We chose to model species-habitat relationships among songbirds that are likely to be either passively targeted for conservation as sagebrush obligates, or influenced by conifer management, and typically appear in conservation planning documents (e.g., Gillihan, 2006). We overlaid past conifer cuts conducted through the Sage Grouse Initiative with predictive distributions of declining songbirds to determine if conifer management for sage grouse has passively targeted or avoided certain species.

## Methods

### Study Area

Our aim was to model species distributions inhabiting the sagebrush (*Artemisia* spp.) ecosystem within the western US. This geography encompasses a diversity of public and private land tenures and jurisdictional boundaries, and is largely defined by cover of both sagebrush- and grassland-dominant understories. Domestic livestock grazing is the primary land use among intact sagebrush steppe, while major anthropogenic factors contributing to habitat loss and fragmentation vary spatially and include infrastructure associated with energy development, cultivation, and urban development. Persistent ecosystem threats also include invasion of exotic annual grasses (e.g. *Bromus tectorum*) and conifer expansion. To best capture a sampling frame representative of sagebrush landscapes, we merged boundaries defined by sagebrush cover with the addition of existing sage grouse Priority Areas for Conservation and management zones (Sage Grouse Initiative, n.d.) and the historic sage grouse species range (Runge et al., 2019; Supplemental Fig. 1).

### Avian Count Data

Selected species were those that are commonly identified in sagebrush and pinyon-juniper management plans including woodland obligates ash-throated flycatcher, gray flycatcher, gray vireo, juniper titmouse, and pinyon jay; and sagebrush-reliant Brewer’s sparrow, green-tailed towhee, sagebrush sparrow (*Artemisiospiza nevadensis*) and sage thrasher (Table 1). We used point count data from the U.S. Geological Survey’s Breeding Bird Survey (BBS), an annual roadside survey conducted from late May - early July by citizen-scientists skilled in avian identification (Pardieck et al., 2017). Along each survey route, participants conduct 50, 3-minute point counts approximately every 0.8km (i.e. ∼40 km routes), and record every bird seen or heard within 400 m. We digitized stop locations using available information on stop descriptions. When stop descriptions were unavailable we generated equidistant points along routes between known stop locations, or between the beginning and ending points of survey routes when no stop locations for routes were available. In total, we had data available for 30,888 stops from 625 BBS routes. We constrained our sample from 2011-2016 such that our response data could best match contemporary spatial predictor variables.

### Spatial Covariates

We broadly hypothesized that heterogeneity in species counts would be influenced by patterns of vegetation, topography, anthropogenic disturbance, fire history, and weather and climate (Table 2). Specifically, we measured the proportion of summarized vegetation types around point counts that were classified as sagebrush, non-sagebrush shrublands, conifer, pinyon-juniper woodlands, scrub- and woodlands, and riparian areas (Table 2). We also combined cropland and developed cover types to characterize anthropogenically disturbed areas. Topography is an important component in structuring bird communities in sagebrush steppe (Knick et al., 2008), so we included measures of elevation, terrain ruggedness (TRI), and a multiscale topographic position index (mTPI) that broadly characterizes landforms (e.g. valley bottoms, ridges, etc.) within 270m, 810m, and 2430m, such that the metric can differentiate between both local- and broad-scale geomorphological features (Theobald et al., 2015). Fire is a pervasive disturbance among sagebrush steppe landscapes structuring vegetation patterns, with the potential for long-lasting negative impacts to densities of breeding birds (Holmes and Robinson, 2013).

**Table 2.** Candidate variables used to describe heterogeneity in Breeding Bird Survey stop level occurrence and count data. Variables were represented as either a mean value across circular spatial windows, or summarized over a temporal window when data was available.

| Variable (abbreviation) | Spatial Window | Temporal Window |
| --- | --- | --- |
| Sagebrush (sage)<sup>a</sup> | 120m, 1km, 6.4km | NA |
| Grassland/Herbaceous (herb) | 120m, 1km, 6.4km | NA |
| Shrubland (shrb)<sup>a</sup> | 120m, 1km, 6.4km | NA |
| Conifer (conf)<sup>a</sup> | 120m, 1km, 6.4km | NA |
| Crop/Disturbed (dist)<sup>a</sup> | 120m, 1km, 6.4km | NA |
| Pinyon/Juniper (piju)<sup>a</sup> | 120m, 1km, 6.4km | NA |
| Scrubland/Woodland (scrub)<sup>a</sup> | 120m, 1km, 6.4km | NA |
| Riparian (ripar)<sup>a</sup> | 120m, 1km, 6.4km | NA |
| Burned Area (burn)<sup>b</sup> | 120m, 1km, 6.4km | 5, 10, 15 years |
| Avg Min Temp (tmin)<sup>c</sup> | 1km | May 15 - Jul 15 |
| Avg Max Temp (tmax)<sup>c</sup> | 1km | May 15 - Jul 15 |
| Total Spring Precip (sprp)<sup>c</sup> | 1km | Mar 15 - Jul 15 |
| Total Winter Precip (wprp)<sup>c</sup> | 1km | Dec 1 - Mar 14 |
| NDVI (ndvi)<sup>d</sup> | 6.4km | May 15 - Jul 15 |
| Elevation (elev)<sup>e</sup> | 30m | NA |
| TRI (tri)<sup>f</sup> | 1km | NA |
| Multiscale TPI (tpi)<sup>g</sup> | NA<sup>1</sup> | NA |
| PDSI (pdsi)<sup>h</sup> | NA | June |

<sup>a</sup> Rollins (2009); <sup>b</sup> Eidenshink (2007); <sup>c</sup> Thornton (2012); <sup>d</sup> Landsat-7 imagery courtesy of the U.S. Geological Survey; <sup>e</sup> Gesch (2002); <sup>f</sup> Riley (1999); <sup>g</sup> Theobald (2015); <sup>h</sup> Abatzoglou (2017).
<sup>1</sup> Index summarized from TPI calculated at 270m, 810m, and 2.43km (Theobald et al., 2015)

Therefore, we used spatial data of fire boundaries and measured the proportion of burned areas 5, 10, and 15 years prior to each point count to characterize the potential legacy effects of fire.

We used weather and climate data that likely influence annual settling patterns of breeding birds. Because precipitation is the primary driver of annual herbaceous growth, we measured total precipitation occurring both over winter (Dec 1 - March 14) and spring (March 15 - July 15) as our study area encompassed ecoregions where precipitation both largely occurs during winter (Great Basin) or spring and early summer (Great Plains). We also summarized patterns of temperature as mean maximum and minimum temperatures over the sampling period (May 15 - July 15) to characterize thermal niches for each species. We used the Normalized Difference Vegetation Index (NDVI) to broadly describe site productivity as NDVI has been correlated with critical life stage requirements for breeding birds (Sweet et al., 2015). We calculated mean NDVI across Landsat scenes over the sampling period, and omitted pixels that were identified as cultivated or woodland in an effort to best characterize the productivity of sagebrush steppe habitats. Drought is a major factor shaping sagebrush steppe systems, so we used Palmer’s Drought Severity Index (PDSI) to identify the spatial and temporal patterns of persistent, long-term drought across the study area. We matched all temporally-referenced weather and climate data to the year of observation across surveys. We also resampled all data to 120m resolution rasters for prediction as spatial covariates varied in their native resolutions (Table 1). Lastly, we included stop (1-50) as a covariate in all models as a proxy for time of day, which is known to influence detection of birds (Niemuth et al. 2017). All continuous covariates were scaled 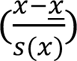 to aid model convergence and coefficient interpretation.

### Spatial and Temporal Scales

We chose three spatial scales to summarize landcover and burned area covariates for consideration in modeling counts including 120 m, 1000 m, and 6400 m. The smallest scale (∼4.5 ha) corresponds to minimum territory sizes of songbirds among sagebrush habitats (Rotenberry et al., 1999). Past work has demonstrated that the amount of shrub and grass cover within 1000 m scale (∼314 ha) has been an important predictor of habitat selection for songbirds in sagebrush steppe (Rotenberry and Knick 1995). This buffer also matches the scale of a typical conifer removal project (e.g. www://conservationefforts.org). Lastly, we hypothesized that a 6400 m scale (∼12,868 ha) represented watershed scale habitat influencing settling patterns by migratory passerines. Past research on sagebrush obligate response to conifer treatment indicated that treatments needed to be adjacent to large intact landscapes (> 14,000 ha) for sagebrush obligate songbirds to recolonize conifer removal areas (Knick et al. 2014).

### Model Fitting, Selection, and Evaluation

Not all species distributions encompassed the entirety of our sampling frame, and we wanted to ensure that we were including only data in analyses that had the potential for a particular species to occur (i.e. 2nd order habitat selection; Johnson, 1980). Therefore, for each species we modeled only data from NACEC level 3 (CEC 1997) that contained >5 detections between 2011-2016, such that predictions are constrained within the occupied range of our sampling frame for each species (Supplemental Fig. 1).

We sought to use one spatial scale to represent each landcover variable, and one spatial and temporal scale to represent a fire variable for each species model. To determine the best fit scales, we fit generalized linear mixed models (glmm) using a binomial error distribution (i.e. detected, undetected) for each landcover and fire variable independently, including random effects for year, route, and BBS observer to account for known sources of heterogeneity in BBS count data (Niemuth et al., 2017). We used detection/non-detection data for scale selection so we wouldn’t have to make assumptions about the proper error distribution for counts, and assumed that the inherent relationship between occurrence and abundance (Royle and Nichols, 2003) would capture relevant variables for count-based models. Using glmm in this step allowed us to calculate Akaike Information Criteria (AIC), from which we used the minimum value among landcover and fire variables to determine the best spatial and temporal scale among variables. Once we had determined a set of candidate variables for each species, we identified highly correlated variables (|r| > 0.6) and removed correlated variables that had lower support determined by higher AIC values.

Once candidate variables for each species were identified, we fit multi-variable models using a random forest approach with regression trees (Breiman, 2001). We modeled count data with random forests regression trees because it was efficiently implemented without making assumptions about an appropriate error distribution and model structure across species, and is generally found to outperform parametric species distribution models in predictive performance (Elith, 2019).

We built regression models using 3000 trees, with a third of the total variables sampled at each split (the default for RF regression; Breiman, 2000). We used fixed categorical effects for years and included latitude and longitude as predictors across models. We evaluated the predictive capability of each species model using k-folds cross validation with 10 folds. For each fold across models we calculated a receiver operator characteristic (ROC) curve and calculated the area under the curve (AUC) by converting predicted counts to measure of occurrence assumed from a Poisson distribution (Royle and Nichols, 2003). We used the mean value of temporally-variant weather and climate predictors and stop number over the study duration, and used 2016-year intercept and fire data to generate spatial predictions for each species.

### Applying Models to Past Conifer Management

Effectively targeted conifer management for both sagebrush and woodland songbirds would take place in areas with higher occurrence of imperiled sagebrush obligates, while avoiding similarly high occurrence areas for declining woodland-dependent species. To test the spatial relationship of past conifer management with songbird occurrences, we evaluated spatial data on conifer management projects from the US Department of Agriculture’s Natural Resources Conservation Service, Sage Grouse Initiative (*hereafter* SGI). As one of the largest restoration efforts in the sagebrush biome since 2010 (Maestas et al. 2021, NRCS 2021), we considered SGI representative of modern conifer removal projects specifically targeted for sage grouse and sagebrush ecosystem restoration. We extracted predicted values for declining species as identified by BBS (Brewer’s sparrow, Sage Thrasher, Green-tailed Towhee, and Pinyon Jay) among the footprint of all SGI conifer treatments (n = 3342; mean = 549.7 ha), and among all the predicted values falling outside of treatment areas. We evaluated targeting of conifer management with logistic regression using treatment as the response variable (1 = treatment, 0 = no treatment), and predicted values for each species as the dependent variable. We reasoned that estimating a positive coefficient would be indicative of conifer management targeting for a particular species; in other words management was taking place in areas of higher predicted occurrence. Conversely, a negative coefficient would imply conifer management was ostensibly avoiding a particular species.

## Results

Our sampling frame encompassed 24 ecoregions (Supplemental Fig. 1), of which focal species were determined to occupy from 7 (Gray Vireo) to 22 (Brewer’s Sparrow) ecoregions within the sampling frame (Table 1, Supplemental Fig. 1). Omitting highly correlated variables, and using model selection to choose among spatial and temporal scales resulted in models with 22 (Sagebrush Sparrow) to 30 (Brewer’s Sparrow) candidate variables describing bird response to topography, weather and climate, landcover, and fire history (Table 3). The spatial and temporal windows selected for landcover and fire variables varied among species, ranging from local (120m) to landscape (6400m) and near (5 yr) and longer-term (15 yr) impacts of fire (Table 3), demonstrating heterogeneous responses by species to landscape features at multiple scales. Across species AUC scores indicated at least good predictive ability across all models (AUC>0.92; Supplemental Table 1). Applying models to spatial grids produced predictive surfaces of occurrence and abundance at landscape scales (Figs. 2-3).

**Table 3.**
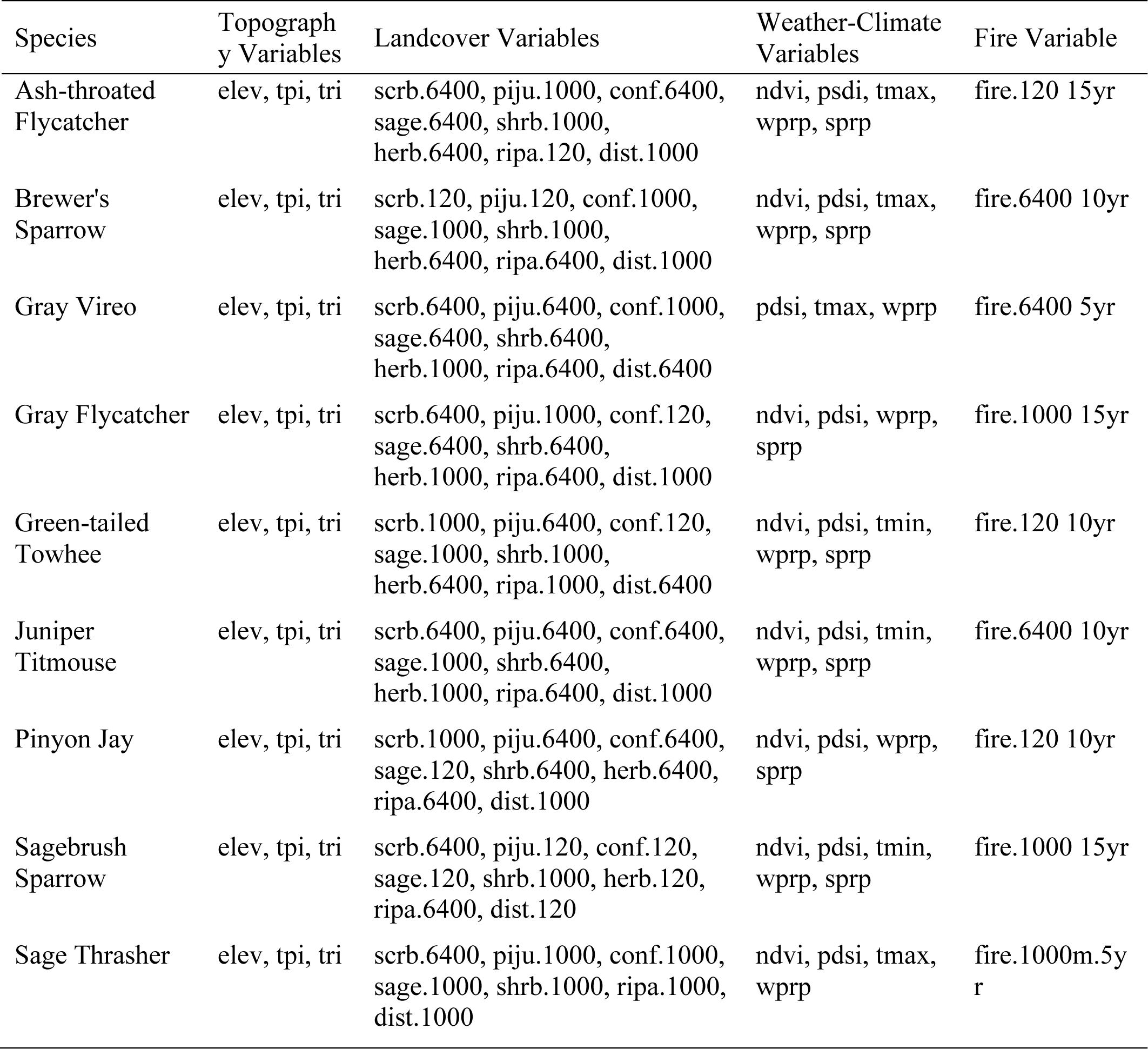
Variables selected for use in models across species with shortened variable codes identified in Table 2. Spatial scales (m) providing the best fit follow the variable name.

**Figure 2.**
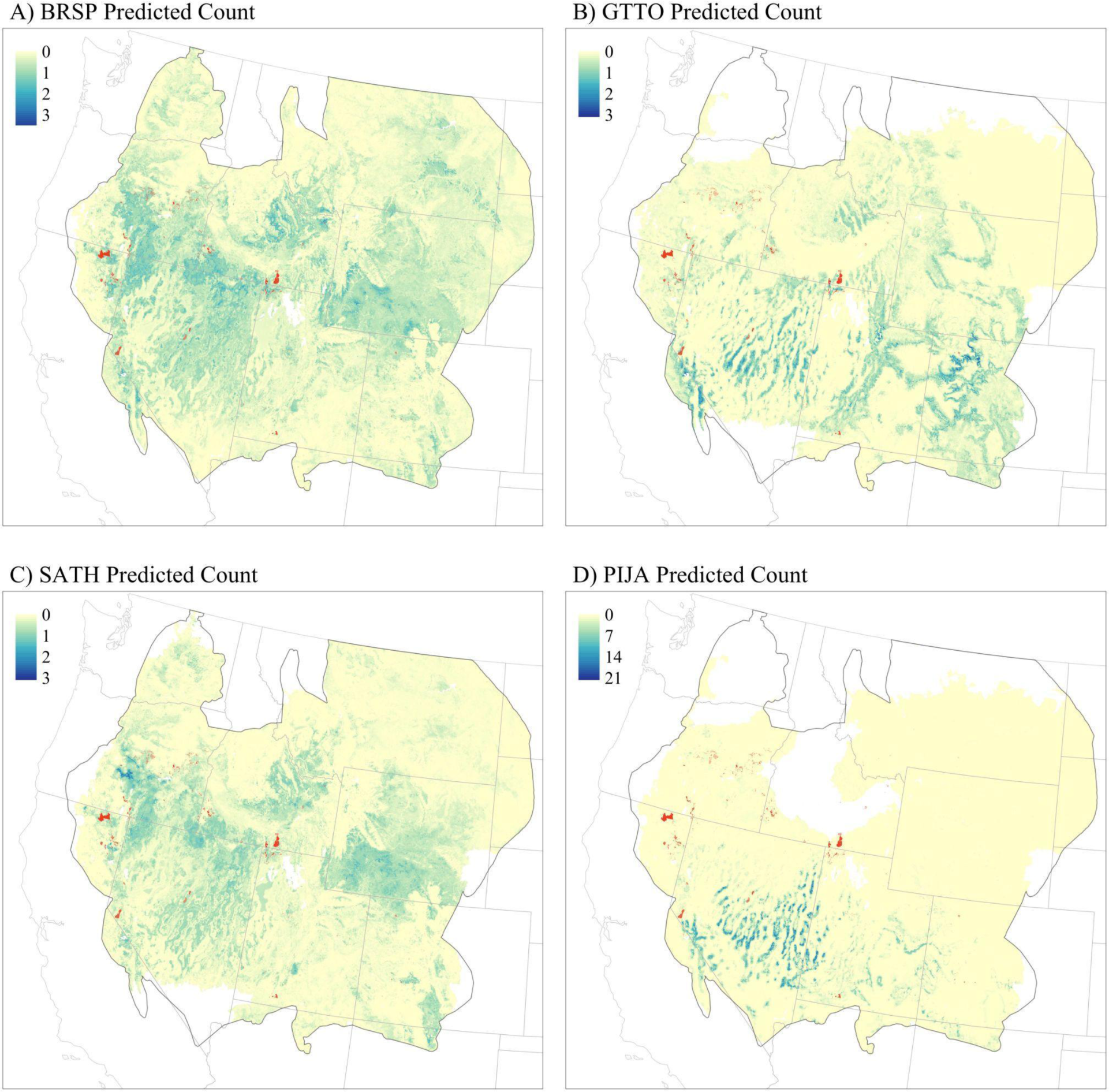
Predicted counts for each of the modeled species with significant declines identified from BBS trends (Table 1) including, A) Brewer’s sparrow (BRSP), and B) green-tailed towhee (GTTO), C) sage thrasher (SATH), and D) pinyon jay (PIJA). Conifer removal projects contracted with the Sage Grouse Initiative are overlaid in red.

**Figure 3.**
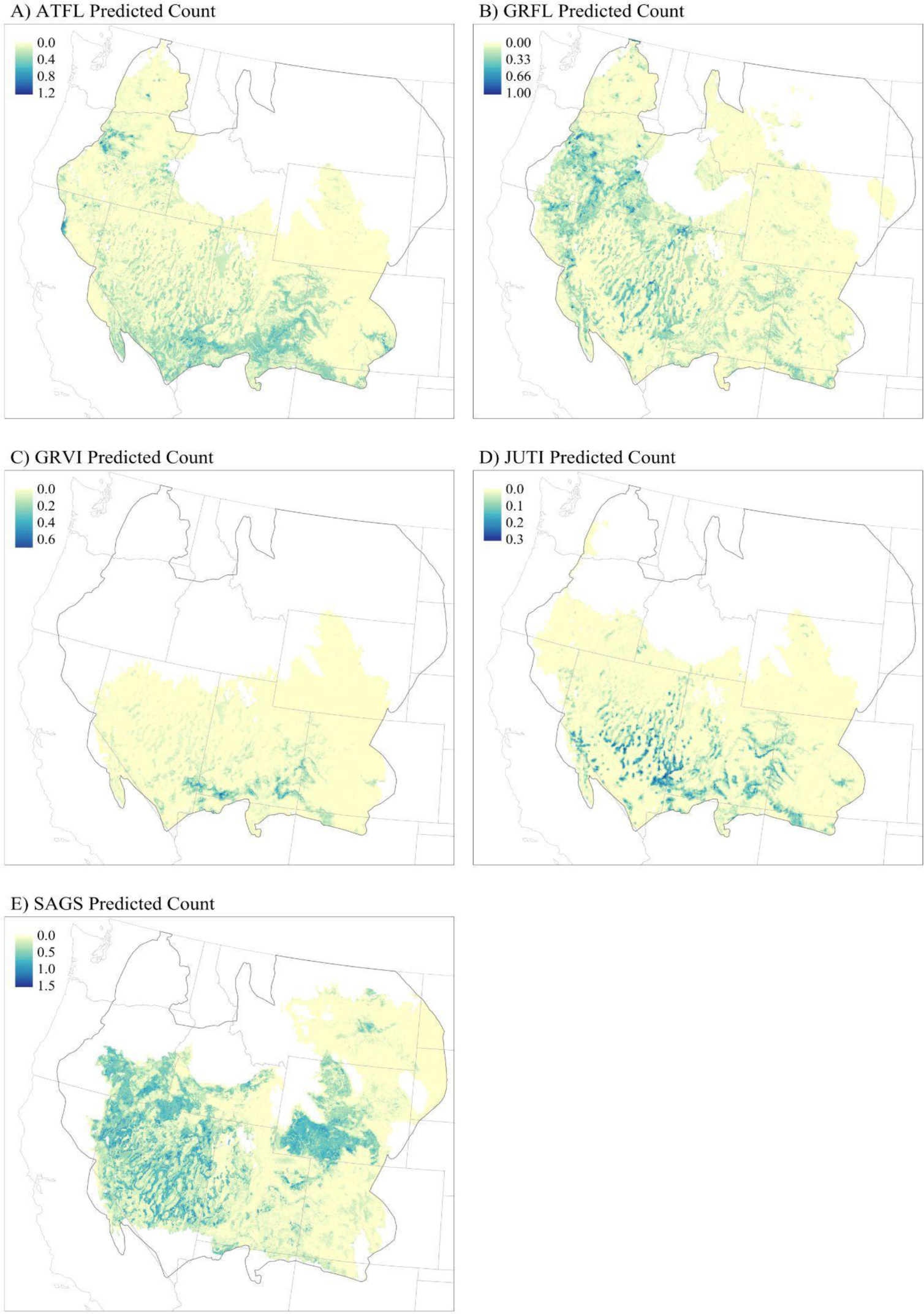
Predicted counts for each of the modeled species with stable to increasing trends identified from BBS trends (Table 1) including, A) ash-throated flycatcher (ATFL), B) gray flycatcher (GRFL), C) gray vireo (grvi), D) juniper titmouse (JUTI), and E) sage sparrow (SAGS).

Logistic regression models fit from overlaying SGI conifer management with models for declining species revealed that past cuts targeted areas with both Brewer’s sparrow (3.468; 95% CI 3.408, 3.529) and sage thrasher (3.12; 95% CI 3.049, 3.198), and avoided pinyon jay (-0.018; 95% CI -0.035, -0.002). Generally, SGI conifer management has been focused in northern distribution of the sagebrush ecosystem including areas in northwest Utah, northern California, and Oregon (Fig. 2).

## Discussion

We provide the first habitat-based maps of songbird distribution and abundance for sagebrush- and woodland-dependent species of high concern across the entire sagebrush biome. These new products expand the spatial targeting toolbox beyond high-profile birds like sage grouse (Doherty et al., 2016, 2010; Row et al., 2018) to empower land managers to incorporate multiple species into holistic conservation strategies. To further aid conservation planning, map-based models are made available for visualization using an online application (https://map.sagegrouseinitiative.com/).

Sage grouse have been identified as an umbrella species for wildlife conservation among sagebrush habitats, an assumption that has been tested with mixed results by measuring the co-occurrence of overlapping species distributions. Generally, distribution and abundance of sage grouse have been found to correspond with other sagebrush-dependent wildlife at regional and biome-level scales (Hanser and Knick, 2011; Pilliod et al., 2020; Rowland et al., 2006; Smith et al., 2019); though results become equivocal for more localized investigations of overlap (Carlisle et al., 2018; Carlisle and Chalfoun, 2020; Smith et al., 2021). Perhaps a more meaningful surrogate measure is to test benefits afforded to multiple species under conservation actions intended to benefit a flagship species such as the sage grouse. We found that conifer removal targeted for sage grouse through SGI also targeted important habitats for declining sagebrush obligate songbirds, a guild of species with a proclivity for positive response to management (Holmes et al., 2017). Community-level benefits from targeted conifer removal is encouraging yet not surprising given similar findings demonstrating high overlap between Brewer’s sparrow and SGI conifer removal (Donnelly et al., 2017), and research demonstrating that landscapes across all US sagebrush steppe habitats targeted for sage grouse conservation (i.e. Priority Areas for Conservation) have been judiciously designed in light of affording protections to sagebrush-reliant wildlife communities (Runge et al., 2019).

Our results also reveal that conifer removal efforts targeted for sage grouse largely avoid areas of high predicted occurrence for pinyon jay. These findings provide the first quantitative assessment demonstrating that targeted sage grouse habitat restoration under one of the largest conservation initiatives in the biome does not appear to be at odds with protecting pinyon jay populations across most of the sagebrush biome despite suggestions to the contrary (Boone et al., 2018; Magee et al., 2019). This disparity is explained in part by the SGI’s private lands emphasis and science-based approach that prioritizes removal of early successional conifer expansion among shrub and herbaceous dominated landscapes (Falkowski et al., 2017; Reinhardt et al., 2017). Colloquially known as “phase 1” woodlands (Miller and Miller, 2007), these areas are characterized by expansion of conifers into shrublands historically devoid of trees, which are used by pinyon jays in some areas mainly for food caching (Boone et al. 2021). Even with the increased attention in sage grouse focused conifer projects over the past decade, the combined effects of management and wildfire are estimated to have only reduced the conifer footprint by 1.6% across the entire sage grouse range (Reinhardt et al. 2020). This pace and scale of removal may barely be keeping up with continued patterns of expansion and infill, which is estimated at 0.4-1.5% annually (Sankey and Germino, 2008).

One area meriting further investigation is the situation in the Central Basin and Range Ecoregion where the highest concentrations of pinyon jay occur, and contemporary pinyon-juniper woodland change is affecting habitat conditions for pinyon jay and sage grouse. In recent decades, pinyon-juniper woodlands in this region have continued to undergo extensive change in stand structure and composition due to increasing conifer densities (Filippelli et al. 2020), resulting in the infill of shrub and tree co-dominant stands (Miller et al., 2008; Romme et al., 2009). Preferred pinyon jay breeding habitat is often described as heterogenous stands of pinyon-juniper and shrubs, that support high cone pine productivity and resulting pinyon nut production (Balda, 2002). Pinyon pine tree vigor, a surrogate for tree productivity, was an important predictor of pinyon jay nest-site selection, which declined with increasing tree size and density (Johnson et al., 2017). Given changing woodland conditions, we hypothesize that pinyon jay habitat use may be shifting to encroached sagebrush shrublands as historic woodland ecological sites become less suitable - an outcome that imperils both pinyon jay and sage grouse which historically occupied different niches and ecological sites along the shrubland-to-woodland continuum (Fig. 1). This hypothesis is consistent with regional population trends showing sagebrush obligate birds and pinyon jay in decline, while species reliant on dense, persistent woodlands have increased (Table 1). Simply avoiding conifer removal projects in occupied pinyon jay habitats is unlikely to be effective with ongoing woodland dynamics, so it may instead be beneficial for managers to consider site-appropriate silvicultural prescriptions designed to restore and maintain the heterogeneous woodland structure critical to pinyon jays.

Co-produced science and monitoring should be coupled with any such restoration efforts (Naugle et al. 2020) to help overcome existing knowledge gaps in woodland restoration (Boone et al. 2018). Evaluation metrics suggested that all species models provided “good” predictive capability across a large geography. However, there are several important caveats when using BBS data to develop spatial models. Notably, BBS sampling occurs along roadways, which could bias the habitats and bird communities observed, and without repeated samples provides only an index to abundance. Past studies within our sampling frame have found no significant differences between counts of sagebrush obligates on road and off-road surveys (Rotenberry and Knick, 1995), nor performance of spatial models applied to data when comparing BBS with samples collected off-roads (Mccarthy et al., 2012). Though it remains possible that the entire covariate space (i.e. niche) was not fully sampled for each species by constricting surveys to road sides (e.g. high elevation roadless sites). Ultimately indices such as relative abundance from BBS data can still provide management with meaningful information on patterns of avian occurrence (Johnson, 2008; Niemuth et al., 2017), particularly when it represents the primary data source available at the temporal and spatial scales relevant to management. Viewed in total, local and design-based studies that employ random sampling and account for detection probability will only improve the quality of spatial planning tools for practitioners, and should be used when available. Similarly, model-based inference at the biome scale will never surpass local knowledge or site evaluation prior to management when it comes to sensitive resources (e.g. location of a particular nesting colony; Johnson et al., 2016).

Management actions to reduce conifer for purposes other than sagebrush ecosystem restoration (e.g. fuels reduction) may not be similarly inconsequential for pinyon jay conservation, particularly when treatments occur among established pinyon-juniper woodlands.

For example, thinning conifers to reduce fire risk in New Mexico, USA left previously-suitable pinyon jay nesting habitat unoccupied following treatment (Johnson et al., 2018) and conifer thinning in Colorado reduced pinyon jay occupancy at local scales (<4 ha), though treatments also resulted increased pinyon jay occupancy at the scale of management (18-117 ha; Magee et al., 2019), highlighting the importance of landscape-level considerations. Both studies documenting impacts to pinyon jay from conifer removal were outside the occupied range of sage grouse (Schroeder et al., 2004). Thus, extending our spatial modeling approach for the pinyon jay distribution beyond the sagebrush biome could better equip conservationists in southwestern ecoregions with important decision-support tools, particularly as these landscapes face additional pressures of drought-induced tree mortality (Clifford et al., 2011; Fair et al., 2018, Shiver et al. 2021).

## Implications

Pinyon-juniper management is often framed as creating “winners” and “losers” among wildlife species (Bombaci and Pejchar, 2016; Zeller et al., 2021), but spatial context of management efforts relative to species populations is often lacking to assess this beyond local project scales. In fact, nuanced analysis reveals that targeted removal of post-settlement era conifer encroachment in sagebrush shrublands may not be at odds with species that rely on conifers for a portion of their life history (Anthony and Sanchez, 2019; Maestas et al., 2019). Model outputs developed here can help inform management for an additional suite of species of concern that lack the spatial tools necessary to avoid potentially detrimental impacts and target limited resources for restoration. Future efforts could combine our species models with spatial data on known ecosystem threats and potential risks to populations to develop holistic, multi-species management plans. For example, combining species models with high resolution vegetation data (Allred et al., 2021; Rigge et al., 2020) can help practitioners make optimal decisions given a bevy of seemingly competing conservation and management interests (Reinhardt et al., 2017; Ricca et al., 2018).

## Acknowledgements

We are grateful to the many volunteers and staff that support the collection and distribution of data for the Breeding Bird Survey. The Western Association of Fish and Wildlife Agencies provided support through the Sagebrush Science Initiative. C. Wiggins identified BBS stop locations, and S. Somershoe, S. Fields, M. Estey, R. Pritchert, and K. Barnes all provided helpful input. N. Niemuth provided invaluable guidance on the framing of the analyses and writing of the manuscript. The findings and conclusions in this article are those of the authors and do not necessarily represent the views of the U.S. Fish and Wildlife Service. The findings and conclusions in this article are those of the authors and should not be construed to represent any official USDA or U.S. Government determination or policy.

---

**Supplemental Table 1.**
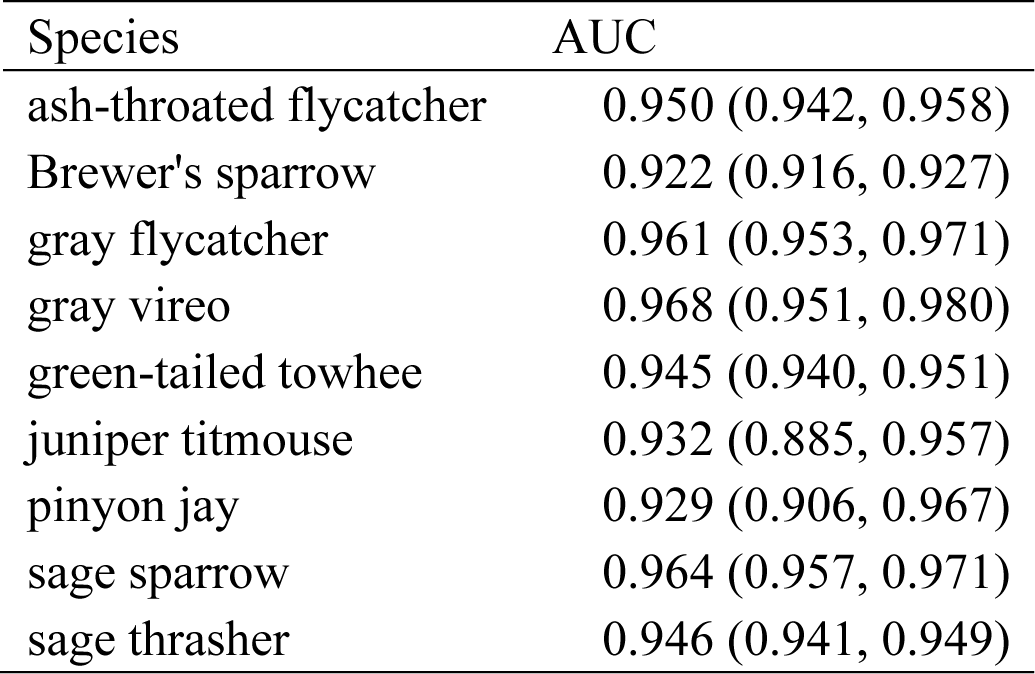
Mean area under the curve (AUC) statistic and range calculated from k-folds cross-validation with 10 folds suggest good model fit across species.

**Supplemental Figure 1.**
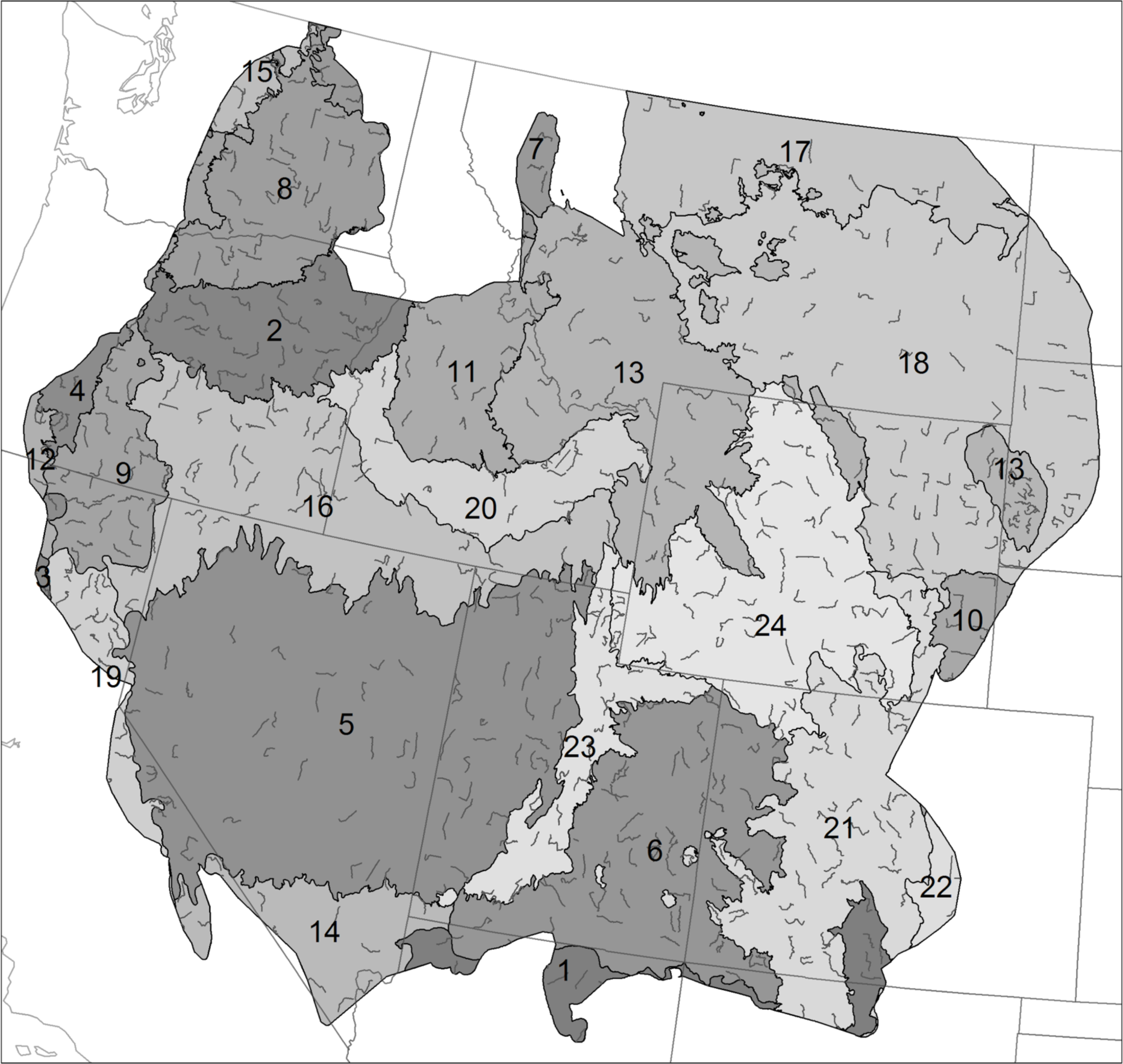
Sampling frame was composed of the US portion of the sagebrush ecosystem as identified by all sagebrush land cover types, with the addition of existing sage grouse Priority Areas for Conservation and management zones (COT 2013), and the historic sage grouse species range (USGS FRESC 2002), which encompassed 625 Breeding Bird Survey routes. We restricted species models Commission for Environmental Cooperation Level 3 ecoregions where they were detected at least 5 times from 2011-2016, which implicated: 1-Arizona/New Mexico Plateau; 2-Blue Mountains; 3-California Coastal Sage; Chaparral, and Oak Woodlands; 4-Cascades; 5-Central Basin and Range; 6-Colorado Plateaus; 7-Columbia Mountains/Northern Rockies; 8-Columbia Plateau; 9-Eastern Cascades Slopes and Foothills; 10-High Plains; 11-Idaho Batholith; 12-Klamath Mountains; 13-Middle Rockies; 14-Mojave Basin and Range; 15-North Cascades; 16-Northern Basin and Range; 17-Northwestern Glaciated Plains; 18-Northwestern Great Plains; 19-Sierra Nevada; 20-Snake River Plain; 21-Southern Rockies; 22-Southwestern Tablelands; 23-Wasatch and Uinta Mountains; and 24-Wyoming Basin.

